# A Comprehensive Roadmap Towards the Generation of an Influenza B Reporter Assay Using a Single DNA Polymerase-Based Cloning of the Reporter RNA Construct

**DOI:** 10.1101/2021.11.19.469260

**Authors:** Nandita Kedia, Saptarshi Banerjee, Arindam Mondal

## Abstract

Mini-genome reporter assay is a key tool for conducting RNA virus research. But, procedural complications and lack of adequate literature pose a major challenge in developing these assay systems. Here, we present a novel yet generic and simple cloning strategy for the construction of influenza B virus reporter RNA template and describe an extensive standardization of the reporter RNP/ polymerase activity assay for monitoring viral RNA synthesis in an infection free setting. Using this assay system, we, for the first time showed the effect of viral protein NS1 and host protein PKC-Delta upon influenza B virus RNA synthesis. Additionally, the assay system showed promising results in evaluating the efficacy of antiviral drugs targeting viral RNA synthesis and virus propagation. Together, this work offers a detailed protocol for standardization of influenza virus mini-genome assay and an excellent tool for screening of host factors and antivirals in a fast, user-friendly and high throughput manner.

## 1 Introduction

First discovered in 1940(1), the influenza B virus has since been causing significant morbidity and mortality in the global population(2). As per the recent surveillance, (seasons 2010–2018) influenza B viruses are responsible for 15-30% of total influenza-like illness, with several complications like fevers, body ache, fatigue and even life-threatening acute respiratory distress syndrome for patients having pre-existing lung diseases(3,4). There are two different lineages of influenza B virus, Victoria and Yamagata. These circulate in the human population with various degrees of predominance in different influenza seasons(5,6). Due to the constant increase in the influenza B virus related infections and limited cross protection offered by influenza B vaccine against both of these lineages, there is a gradual transition from trivalent (against two subtypes of Flu A and one lineage of Flu B) to quadrivalent (against two subtypes of Flu A and two lineages of Flu B) flu shots offered across the world(7). In spite of its immense importance in the context of global healthcare ecosystem, influenza B virus research has drawn significantly lesser attention compared to the closely related influenza A viruses, largely due to the scarcity of the tools required to study virus replication cycle. This also severely restricts antiviral drug discovery directed towards influenza B virus therapy(8).

Influenza viruses are segmented negative-strand RNA viruses of the Orthomyxoviridae family. Amongst the four types A, B, C and D, only influenza A and B cause human epidemics(9). The viral genome consists of eight different segments each of which remains enwrapped with multiple copies of nucleoprotein (NP) in their oligomeric form and associates with a single copy of RNA dependent RNA polymerase (RdRp) to form the ribonucleoprotein complexes or RNPs(10). RNPs are the self-sufficient machinery for driving different modes of RNA dependent RNA synthesis events including viral gene expression and genome replication, hence reside at the center of virus replication cycle(11). This is why reporter RNP based assay systems remain one of the invaluable tools for studying virus replication, host-pathogen interaction and high-throughput screening of antivirals without handling infectious virus particles and hence avoiding biosafety associated procedural complications(12,13).

Influenza virus genomic segments are single-stranded RNA that are devoid of the 5′-Caps and 3′-Poly(A) tails (14). Different segments harbor conserved untranslated regions of variable lengths both at the 5′ and 3′ ends, which bracket single or multiple open reading frames (in the antisense orientation) encoding viral proteins(14,15). Terminal regions of the 5′ and 3′ UTRs contain complementarity resulting in a partial duplex structure (also known as “panhandle RNA” or “cork-screw RNA”) that serves as the promoter for viral RdRp(16). Additionally, the UTRs contain cis-acting elements, necessary and sufficient for the transcription and replication of viral(17–19) and non-viral reporter genes(16,20–22). Several groups have established reporter RNA based assay systems where viral open reading frames have been replaced with reporter genes of fluorescent, bioluminescent or chemiluminescent proteins(12,13,16,23–26). These reporter RNA templates, when expressed inside the cells in combination with NP and RdRp proteins, reconstitute reporter RNPs. RNA synthesis activity of these RNPs could be measured by quantifying the extent of reporter gene expression. Although this appears a straightforward procedure, but the successful establishment of a reporter assay system requires (i) complicated cloning strategies to synthesize the reporter RNA construct, (ii) construction of plasmids for expression of viral NP and RdRp subunits (PB1, PB2 and PA), and (iii) optimized expression of the reporter RNA and viral proteins in required stoichiometric amounts that leads to the reconstruction of reporter RNPs with maximum efficiency.

So far, different strategies have been used to construct plasmids expressing influenza A and B virus reporter RNA template (reporter plasmid)(12,13,25–27). In a few studies, the reporter luciferase gene was amplified using primers containing long overhangs corresponding to 3′ and 5′ UTR regions of influenza A or B viruses; then, the resulting PCR fragment harboring luciferase ORF flanked by the viral UTRs were inserted into the target vector for RNA polymerase-I driven expression of the same, using conventional restriction digestion and subsequent ligation method(12,22,28). Alternatively, viral 5′ and 3′-UTR containing vectors (amplified using inverse PCR from the cDNA clone of the corresponding segment) were ligated with reporter gene insert predigested with compatible restriction enzyme sites(29). In another cloning strategy, a double-stranded DNA linker encompassing 5′ and 3′-UTRs was inserted into the vector in-between the Pol-I promoter and terminator sequences with the help of compatible restriction enzyme sites. The reporter gene was then inserted between the UTRs using a second restriction enzyme site(25). All of these restriction enzyme based cloning strategies are laborious and often introduce additional nucleotides between the UTRs and the reporter gene, which may interfere with the activity of the cis/trans acting elements(25). In order to avoid these constraints, restriction enzyme free cloning methods utilizing vectors and inserts containing overlapping sequences have also been implemented. For example, inserts containing reporter genes flanked by the 5′ and 3′-UTRs were created using long overhang primers (containing the UTR regions) which was then stitched to the vector through the use of specialized proprietary enzymes/kits(13,30). With the inherent limitations of the aforesaid cloning techniques, scarcity of information about the extensive experimental protocol makes it difficult to establish and standardize the reporter based RNP activity assay for influenza viruses. This situation gets further complicated for influenza B viruses due to the larger size of the UTRs compared to the same for influenza A viruses.

Here, we present a novel yet fairly simple cloning strategy, independent of any restriction enzyme or specialized reagents or kits, to construct a firefly luciferase based reporter plasmid capable of generating a reporter genome template for influenza B/Brisbane/60/2008 virus. Additionally, we present extensive standardization of this reporter plasmid-based RNP activity assay through optimization of various parameters regulating viral RNA synthesis. Using the reporter assay system, we have shown for the first time the effect of viral non-structural protein-1 (NS1) and host protein kinase C delta (PKCD) upon influenza B virus RNA synthesis. We have also demonstrated the ability of this assay system to be used as a high throughput screening platform for the identification of antiviral drugs specifically inhibiting RNA polymerase activity of the virus. Together, this work presents a great resource for cloning, standardization and implementation of reporter-based RNP activity assay for influenza and other related viruses.

## 2. Materials and methods

### 2.1 Cell lines and Viruses

Human embryonic kidney 293T (HEK 293T) cells were maintained in Dulbecco’s modified Eagle’s medium (DMEM; Gibco™, Cat no. #12800017) supplemented with 10% (v/v) fetal bovine serum (FBS; Gibco™, Cat no. #10082147), 2mM GlutaMAX™ (Gibco™, Cat no. #35050061), 1% penicillin-streptomycin (Gibco™, Cat no. #1514122), incubated at 37°C in a humidified 5% CO2 incubator. Madin-Darby Canine Kidney (MDCK) cells were maintained in the same conditions with 10% FBS (Gibco™, FBS; Cat no. #10270106). Influenza B/Brisbane/60/2008 virus was used in this study.

### 2.2 Virus amplification and RNA Extraction

3×10^6^ MDCK cells were seeded in 10cm dishes, 24 hours before the infection. Prior to infection, the cell monolayer was washed with PBS twice and subsequently infected at an M.O.I. of 0.001. For each 10 cm dish, 1 ml virus inoculum was prepared in virus growth media (VGM; containing DMEM, 0.2% bovine serum albumin (Sigma; Cat no. #A8412), 25 mM N-(2-hydroxyethyl) piperazine-N′-ethanesulfonic acid (HEPES; Invitrogen, Cat no. #15630080) buffer, 2 mM GlutaMAX™, 1% penicillin-streptomycin and 0.5 µg/ml TPCK-trypsin (Thermo Scientific™, Cat no. #20233)). Virus attachment was performed with 1ml of inoculum for 1 hour at 37 °C in humidified 5% CO_2_ incubator with intermittent shaking at every 10 minutes to prevent drying of cell monolayer and homogenous distribution of the inoculum. Post attachment, each 10 cm dish was supplemented with 6 ml of VGM and incubated at 33 °C in a humidified 5% CO_2_ incubator. At 72 hours post-infection, the supernatants were collected and centrifuged at 3200 g for 10 minutes at 4 °C to remove the cell debris. The supernatants were collected and aliquots were stored at 80 °C refrigerator for further applications(31).

### 2.3 Reverse transcription (RT)-PCR

Viral RNA was extracted from amplified virus stock using Trizol reagent (Invitrogen; Cat no. #15596018). Reverse transcription was carried out using in-house produced Moloney murine leukemia virus (MMLV) Reverse transcriptase (RT) enzyme and the primer ‘Uni9’, which is complementary to all the segments of Influenza B/Brisbane/60/2008 virus. Sequences of primers are mentioned in Table 1. Briefly, 10 µl of viral RNA (∼500 ng), 1 µl of 2 µM Uni9 primer and 1µl of 10 mM dNTP (Thermo Scientific™, Cat no. # R0181) were mixed and warmed at 65 ºC for 5 minutes. This is followed by snap chilling at ice for 2 minutes. After snap chilling the reaction, the premixed solution containing 4 µl of 5×RT buffer (Invitrogen, Cat no. #18080044), 2µl of 0.1 M DTT (kit component of Invitrogen, Cat no. #18080044), 0.25 µl of RNase inhibitor (Thermo Scientific™, Cat no. #EO0381), 1µl of MMLV RT and 0.75 µl of sterile nuclease-free water (AMRESCO, Cat no. # E476-1L) were added to the tube containing snap chilled mix. The reaction was carried out at 42ºC for 50 minutes and was terminated by heating at 70 ºC for 15 minutes. After the RT reaction, in which cDNA for all 8 segments have been generated; the PCR reaction was carried out using NA-NB segment specific primer to enrich the fragment specific for segment 6 using the primers ‘NA-NB_F’ and ‘NA-NB_R’(primer sequences are listed in Table 1). Each of the 50 μl PCR reaction mixture consisted of 10 μl of 5×Phusion HF Buffer, 5 μl each of 5 μM primers, 5 μl of 2 mM dNTPs, 5 μl each of the synthesized cDNAs, 19.5 μl of sterile nuclease-free water, and 0.5 μl of Phusion High-Fidelity DNA Polymerase (Thermo Scientific™, Cat no. #F530S). The cycling conditions for the PCR reaction is stated in Table 2. For RT reactions, the lyophilized primers were dissolved and diluted in sterile nuclease free water. For PCR specific primers, the initial stocks were dissolved using 10 mM Tris-HCl (pH=8) and working stocks were diluted in sterile nuclease free water.The PCR products were purified using PCR Purification Kit (Invitrogen; Cat No. # K310001). The yield and quality of the purified products were checked measuring the UV absorbance at 260 nm and 280 nm, and subsequently by running it on agarose gel electrophoresis.

**Table 1.**
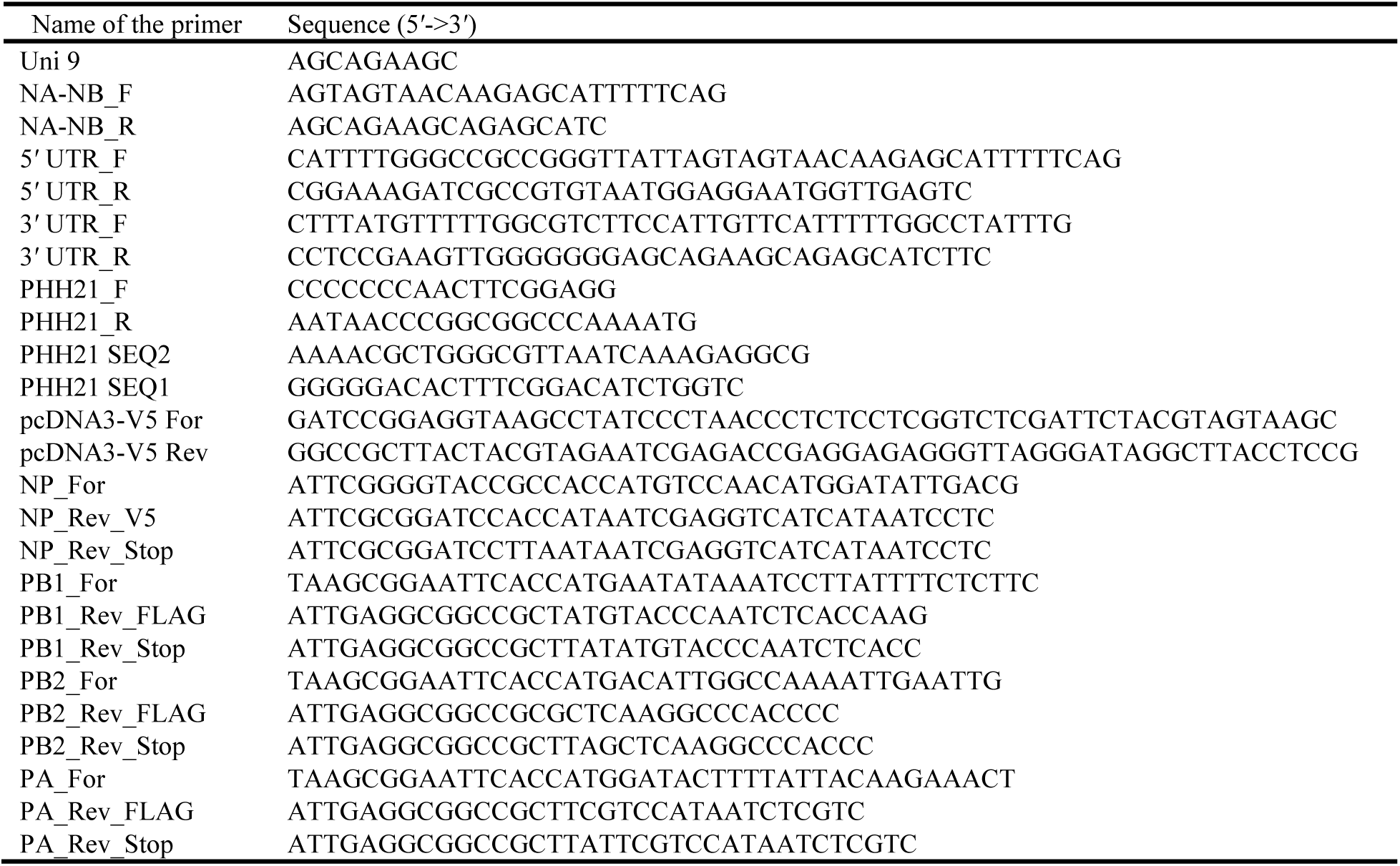
Primers used in this study.

**Table 2:**
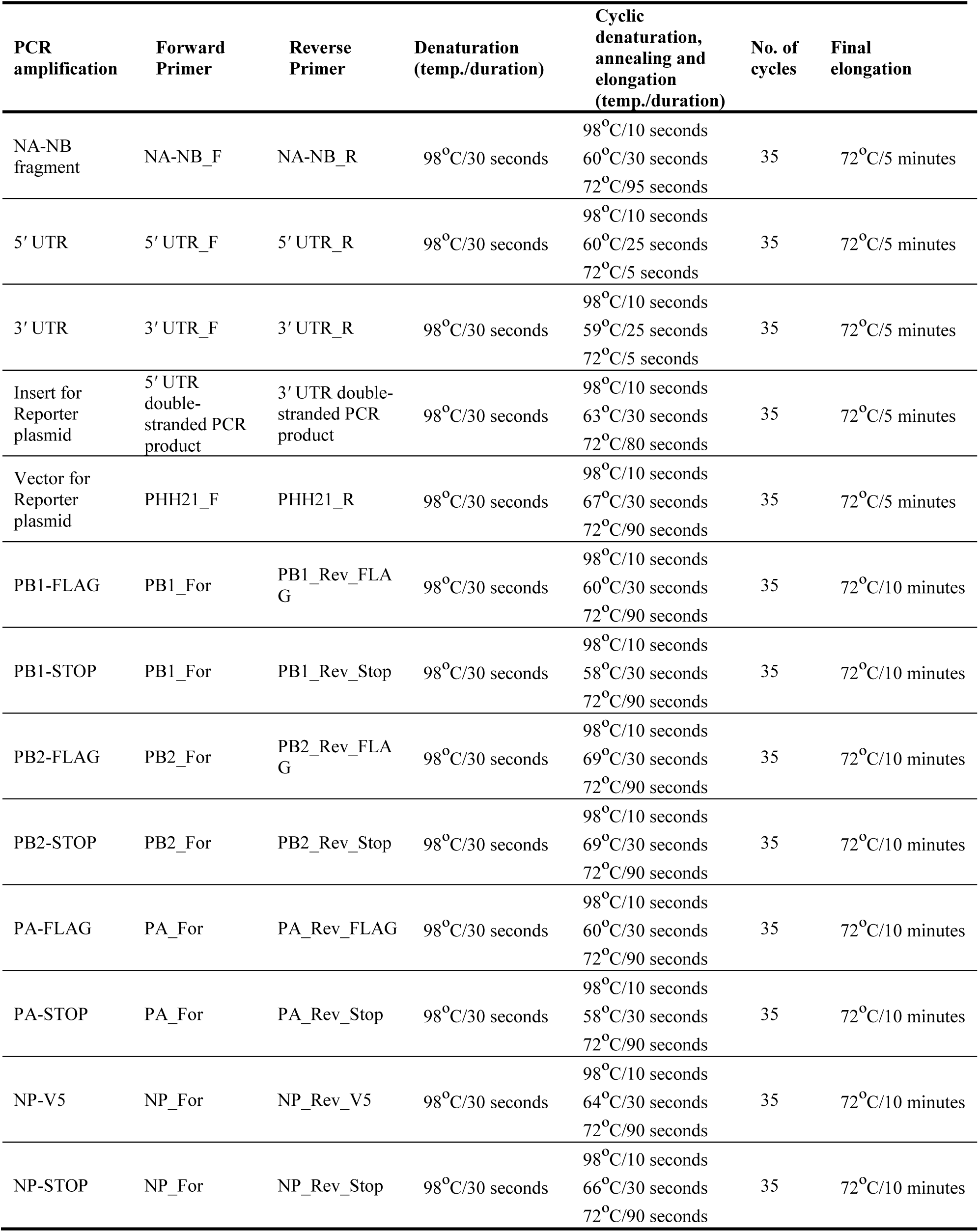
PCR conditions used in the study for amplification of individual inserts.

### 2.4 PCR and Cloning

#### 2.4.1 Amplification of 5′ and 3′ UTR

The RT-PCR amplified DNA corresponding to segment 6 of the viral genome was used as a template for the amplification of 5**′**UTR and 3**′**UTR using Phusion High-Fidelity DNA Polymerase using 5′ UTR_F, 5′ UTR_R and 3′ UTR_F, 3′ UTR_R primers pairs, (all the primer sequences are listed in Table 1 and the PCR conditions are listed in Table 2) respectively.The PCR product were purified using Quick gel extraction kit (Invitrogen: Cat no. # K210012). The purified double-stranded 5′UTR fragment and 3′UTR fragment (containing overhangs for insert and vector fragment) were then used as primers for the amplification of luciferase gene (insert amplification) as described in the next section.

#### 2.4.2 Preparation of Insert

The luciferase ORF was amplified using the pHH21-vNA-Luc as a template, kindly provided by Dr. Andrew Mehle, as a template. The plasmid encodes firefly luciferase gene flanked by Influenza A UTR’s. This plasmid encodes firefly luciferase gene flanked by Influenza A UTR’s. The double-stranded 5′UTR and 3′UTR fragments, synthesized in the previous step, were used as primers (5µM final concentration) for the PCR amplification of Luciferase ORF using Phusion high fidelity DNA polymerase, following manufacturer’s protocol (for primer sequences and PCR conditions, refer to Table 1 and Table 2, respectively). The PCR product was analysed on 0.8% agarose gel and purified using PCR purification kit.

#### 2.4.3 Preparation of Vector

The vector was amplified using the pHH21-vNA-Luc as a template. PHH21_F’ & ‘PHH21_R’ primers (listed in Table 1) were used to amplify and linearize the vector using Phusion high fidelity DNA polymerase using 5x Phusion GC rich buffer following manufacturers’ protocol. Primer sequences and PCR conditions are listed in Table 1 and Table 2, respectively. The PCR product was analysed on 0.8% agarose gel and purified using PCR purification kit.

#### 2.4.4 Circular Polymerase Extension Cloning

In the final CPEC assembly and cloning reaction(32), four different CPEC reactions have been set up using the purified linearized vector (mentioned in section 3.4.3.) and inserts(mentioned in section 3.4.2.) maintaining a molar ratio (V:I) = 1:0, 1:1, 1:2 and 1:3 respectively(Figure 2D).The reaction mixture composition are described below:

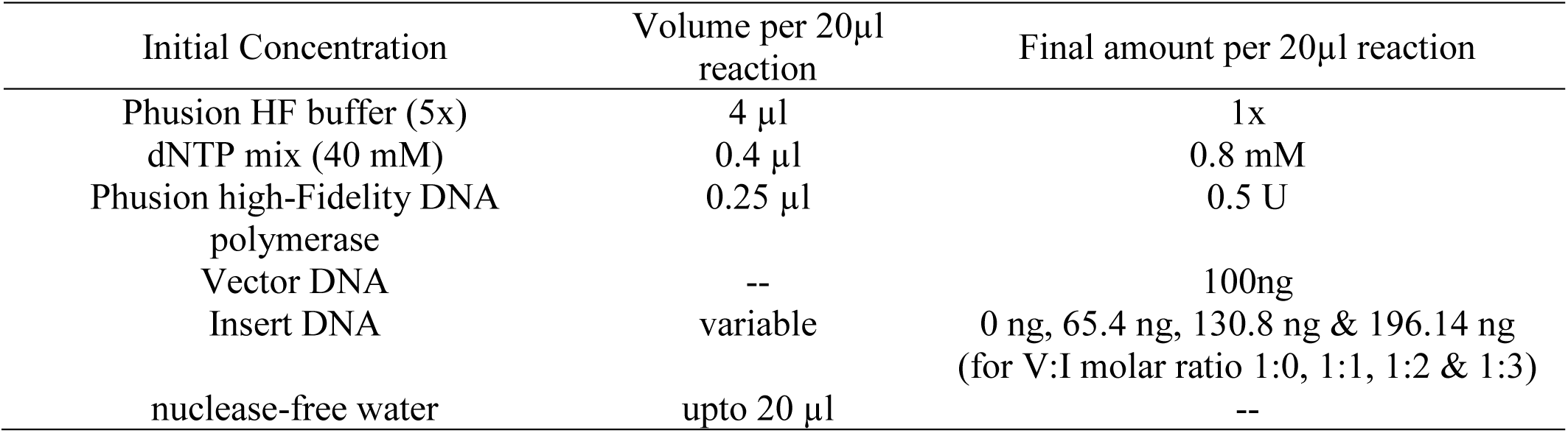

**Figure 1:**
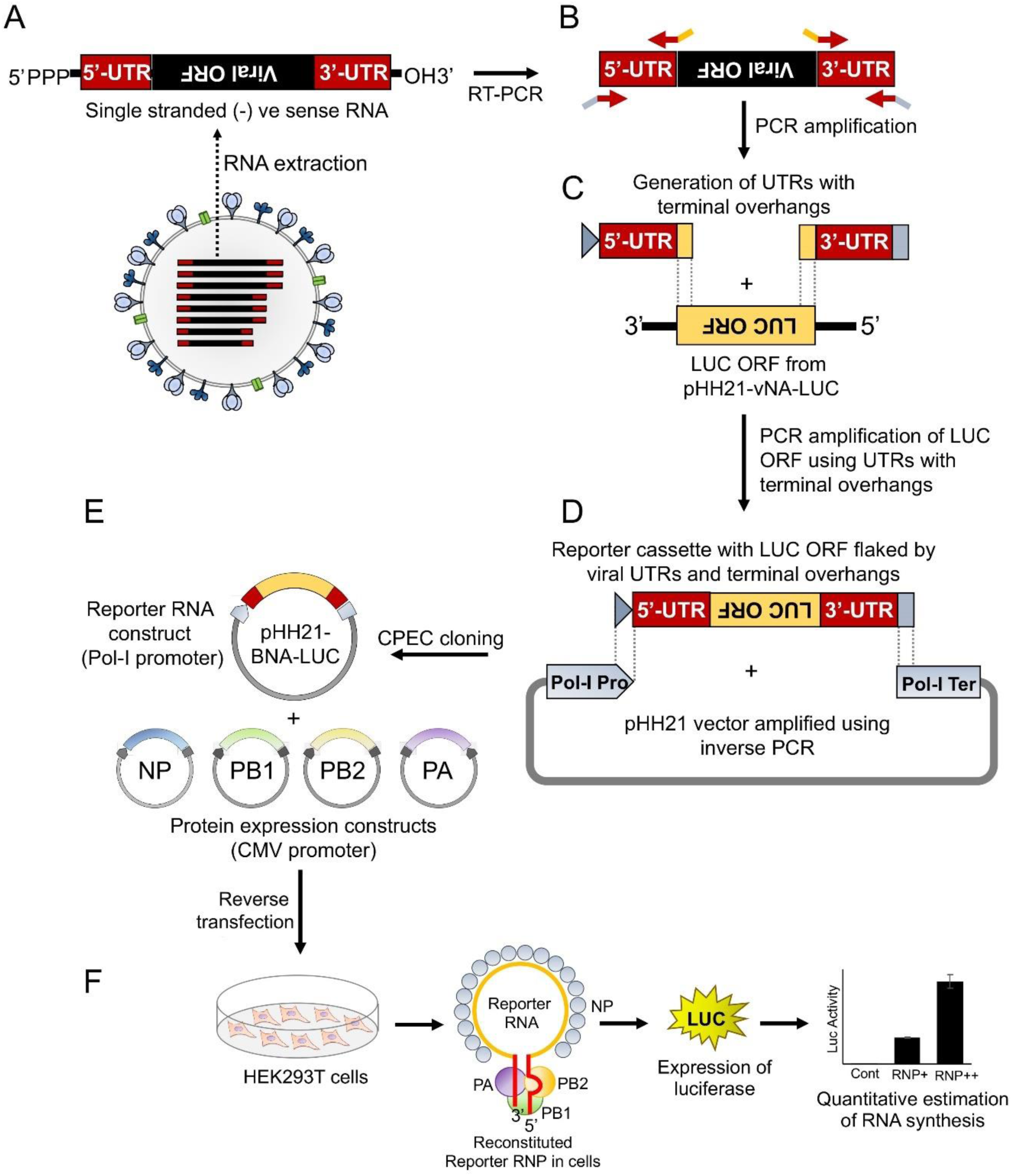
An overview of cloning strategy of influenza B reporter plasmid and reporter assay (A) Total RNA was isolated from amplified stocks of Influenza B/Brisbane/60/2008 virus (B) total RNA was converted to cDNA by performing RT-PCR and 5**′** & 3**′** UTRs were amplified using specific primers containing overhangs. (C) The double-stranded 5**′** and 3**′** UTRs containing overlapping regions were used as primers to amplify the luciferase ORF. (D) The resulting PCR product was used as an insert for CPEC assembly with the PCR amplified vector fragment. (E-F) The generated reporter and other protein expressing plasmids upon co-transfection in HEK293T cells reconstitute the luciferase RNP’s that express luciferase enzyme under the control of the viral promoter. The quantification of the luciferase signal gives the measure of viral polymerase activity.

**Figure 2:**
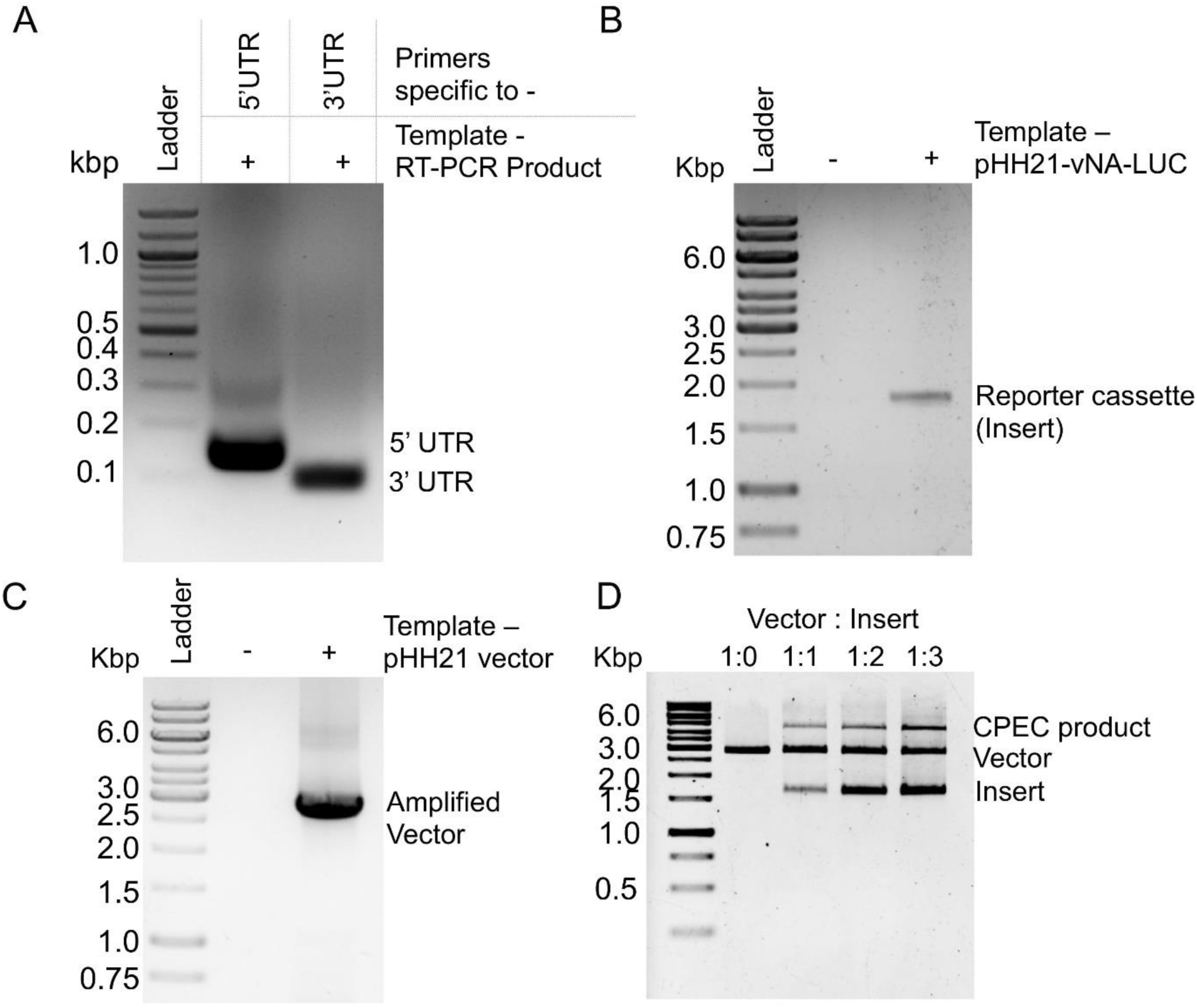
PCR amplification and CPEC reaction for construction of reporter construct: Agarose gel electrophoresis images of (A) PCR amplification products corresponding to the 5**′**UTR and 3′UTR of NA-NB segment. (B) PCR amplification product of luciferase ORF using double-stranded PCR products corresponding to 5**′** and 3**′** UTRs as primers. (C) PCR amplification of pHH21 vector. (D) CPEC products with different ratio of vector to insert.

Cycling conditions for the CPEC assembly and cloning reactions are described below:

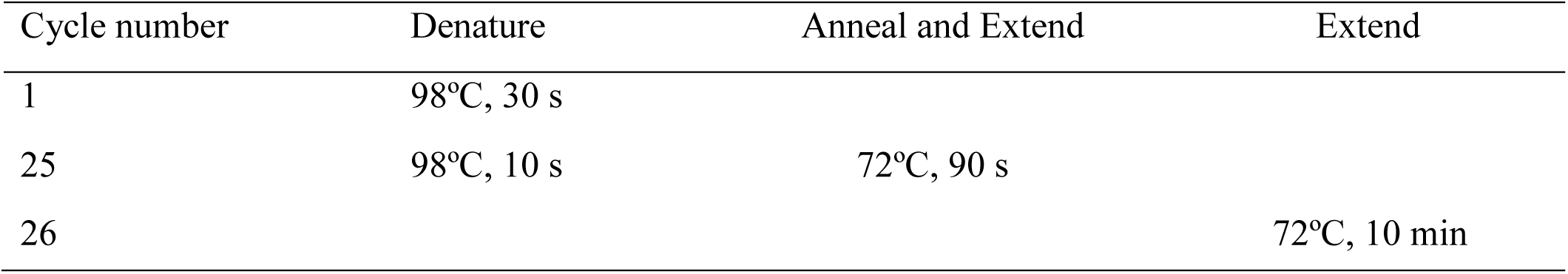

To assess if a CPEC reaction is successful or not, 10µl of each product was analysed on agarose gel electrophoresis (Figure 2D). The V:I =1:3 reaction showing highest intensity of the high molecular weight band (corresponding to the total length of vector and insert) was selected for transformation.

### 2.5 Transformation

*E. Coli*. DH5α competent cells were prepared by modified rubidium chloride method as described by Glover et al.(33). For transformation, 10 µl of the CPEC reaction mixture were added to the competent cells. Followed by an incubation period on ice for 30 minutes, cells were subjected to a brief heat shock at 42ºC for 35 seconds followed by 5 minutes incubation in ice. 400µl of Luria bertani broth (Himedia, Cat no. # M1245; 2.5% in double distilled water) were added immediately after the incubation and the cells were grown for 1.5 hours in 37 ºC incubator with shaking at 220 rpm. Once the shaking period is over, and the entire culture volume was spread upon LB-agar plates containing 100 µg/ml of Ampicillin. Positive clones were identified by colony PCR and confirmed by Sanger’s sequencing.

### 2.6 Generation of polymerase protein expressing plasmids

The PB2, PB1 and PA ORFs were cloned into the pCDNA-3×-FLAG vector (generously provided by Dr. Andrew Mehle) which is a modified version of pcDNA3.1 (addgene) vector expressing proteins under CMV promoter. This vector contains three FLAG epitopes joined in tandem (3×-FLAG) after the NotI site at its MCS followed by a Cytosine. This results in the expression of a protein having tri-alanine linker in between the individual ORFs and the C terminal 3×-FLAG tag. For the expression of untagged version of each RdRp subunit, the stop codon has been kept intact at the end of ORF. For the expression of V5 tagged NP, pcDNA3 vector has been modified in order to have glycine-glycine-serine-glycine linker in between the ORF and the C terminal V5 epitope tag. Briefly, two primers of 58 nucleotide length (‘pcDNA3-V5 For’ & ‘pcDNA3-V5 Rev’) were annealed to create double-stranded piece of DNA (the V5 linker) having sticky ends on both the sides (BamHI restriction site at the beginning of the sequence and the NotI site at the end of the sequence). The thermal protocol for ramp down annealing was as follows: 95 ºC for 5 minutes, 70 cycles of 95 ºC (-1 ^º^C/cycle) each for 1 minute, followed by hold at 4 ºC. The V5 linker was phosphorylated at 5′ end by treatment with T4 Polynucleotide Kinase (PNK, Cat no. # M0201S). The pcDNA3.1 (addgene) vector was digested with BamHI and NotI, treated with Calf alkaline phosphatase (CIP; NEB, Cat no. #M0290S) and ligated with the V5 linker in V: I = 1:20 ratio.

Each individual insert fragment has been amplified using the cDNA template with the primers containing restriction enzyme overhangs. The PB2, PB1 and PA have been amplified with EcoRI and NotI overhang in two different PCR sets, one omitting the stop codon and the other including the stop codon in the reverse primer. The NP has been amplified using primers with KpnI and BamHI overhangs. For each amplification, 50 μl of PCR reaction consisted of 10 μl of 5×Phusion HF buffer, 5 μl each of the forward and reverse 5 μM primers, 5 μl of 2 mM dNTPs, 5 μl of cDNA template, 19.5 μl sterile nuclease free water and 0.5 μl of Phusion High-Fidelity DNA Polymerase. All the primer sequences and the specific PCR conditions are listed in are listed in Table 1 and Table 2, respectively). The modified pcDNA3-3×-FLAG have been digested with EcoRI & NotI (NEB) and the pcDNA3-V5 vector has been digested with KpnI & BamHI, followed by treatment with CIP. The digested vectors as well as insert fragments were gel excised and ligated in a vector to insert ratio of 1:3 using T4 DNA ligase (Thermo Scientific™, Cat no. # EL0011) as per manufacturer’s protocol. 10 µl of ligation mixture was transformed into DH5α competent cells (as described in section 3.5, except that 150 µl of culture volume was spread upon LB-agar plates at the end of the transformation). All the positive clones were identified by colony PCR and confirmed by Sanger’s sequencing.

### 2.7 Transfection

For examining the protein expression level of each plasmid, HEK293T cells were transfected using lipofectamine 3000 (Invitrogen: Cat no. # L3000015). Each plasmid wasprepared using plasmid DNA isolation kit (Promega, Cat No. # A1222) and 100 ng working stocks were prepared. The pcDNA3.1 blank vector was used for control sets. Briefly, 26µl mixture of optiMEM (25µl, Thermo Scientific™, Cat no. #31985-070) and p3000(1µl) were added to 500ng of plasmid DNA and mixed well. 26.5µl mixture of optiMEM (25µl) and lipofectamine 3000 (1.5µl) were further added to the DNA-p3000 premix, mixed well and incubated for 15 minutes at room temperature. 2.5×10^5^ HEK293T cells were seeded into 24 well plates, the transfection mixture was added to the respective wells and kept for incubation at 37 ºC humidified CO_2_ incubator. The media was changed 12 hours post-transfection and incubated for 36 (or stated otherwise) hours following transfection.

### 2.8 Western Blot

Protein levels for transiently transfected cells were assessed by Western blotting. Transfected cells were lysed for 20 minutes in pre-chilled Co-Immunoprecipitation buffer (50mM Tris-HCl pH 7.4, 150mM NaCl, 0.53% NP-40) supplemented with 1×protease inhibitor (Roche-Sigma, Cat no. # 11873580001) and 1×phosphatase inhibitor (Thermo Scientific™, Cat No. #78420). The lysates were centrifuged at 21,000g at 4 ºC to remove the cell debris, the supernatant were collected, mixed with sodium dodecyl sulfate-polyacrylamide gel electrophoresis sample buffer and boiled for 10 minutes. The total protein were separated via 8% SDS-PAGE and transferred to polyvinylidene difluoride (PVDF, Bio-Rad, Cat no. #1620177) membrane using transfer buffer (25 mM Tris, 191 mM glycine, 0.025% SDS & 10% methanol (vol/vol) in Trans-Blot Turbo Transfer System (Bio-Rad). After incubation with 5% nonfat milk in TBST (50 mM Tris, pH 7.4, 150 mM NaCl, 0.5% Tween 20) for 60 min, the membrane was washed once with TBST and incubated with antibodies against FLAG (1:5000, Sigma, Cat no.# F3165), V5 (1:5000, CST, Cat no. # D3H8Q), HA (1:5000, CST, Cat no. # C29F4), GAPDH (1:5000, Biobharati LifeScience Pvt. Ltd, Cat no.# BB-AB0060) BNP (1:5000, generated in collaboration with BioBharati LifeScience Pvt. Ltd, India), at 4°C for 12 h. Membranes were washed with TBST three times for 5 minutes and incubated with a 1:25,000 dilution of horseradish peroxidase-conjugated anti-mouse (Sigma, Cat no. # A9044-2ML6) or anti-rabbit (Sigma, Cat no. # A0545-1ML) antibodies for 1 h. Blots were washed with TBST three times for 5 minutes and developed with the ECL system (ThermoFisher, Cat No. #34095) according to the manufacturer’s protocols.

### 2.9 Polymerase activity assay

For polymerase activity assay, the sets where no additional plasmid DNA were used other than the core RNP components (e.g. Figure 3 D,E, Figure 5 C,D), 2.5×10^5^ HEK293T cells were co-transfected with 118 ng of each of the pcDNA3-PB2-FLAG, pcDNA-PB1, 15 ng of pcDNA-PA, 125 ng of pcDNA3-BNP and 125 ng of pHH21-BNA-Luc plasmids to reconstitute 500ng of total DNA(1/4^th^ BNP, 1/4^th^ pHH21-BNA-Luc, and the half of the total amount of DNA have been divided as 8:8:1 ratio for PB2, PB1 and PA). For the negative control set, pcDNA3.1 blank vector was used instead of PB2 subunit. For the experiment of Figure 3A, 100ng of each polymerase subunit have been transfected which has been updated in the subsequent experiments. For polymerase activity assay with additional plasmid components (e.g. Figure 4 A & B), the total RNP reconstituting plasmids have been reduced to 400 ng by transfecting 94.11 ng of each of the pcDNA3-PB2-FLAG, pcDNA-PB1, 11.76 ng of pcDNA-PA, 100 ng of pcDNA3-BNP and 100 ng of pHH21-BNA-Luc expressing plasmids. The additional plasmids were used in various amounts (NS1:50ng & 75ng; PKC delta Cat:15ng, 30ng, 60ng & 90ng respectively) and topped up by blank vector upto 100 ng in order to keep the total amount of DNA same in all the sets.. The transfection mix were prepared with Lipofectamine 3000 as stated earlier. All the transfections for luciferase activity assay have been performed in triplicates. At 12 hours, the media was changed very carefully without dislodging any cell to avoid manual variation among the sets. The cells were harvested at 36 hours (or as mentioned) post transfection and luciferase activity assay was performed using Promega Luciferase Assay System (Promega, #E1500 & #E1910). Briefly, the media was aspirated and 250µl (1x CCLR for firefly luciferase assay) & 100 µl (1x PLB for dual luciferase assay) of lysis buffer (supplemented with 1x protease inhibitor and 1x phosphatase inhibitor) was added to each well and incubated for 20 minutes at 4 ^º^C (firefly luciferase assay) and room temperature (dual luciferase assay) respectively. The assay was performed as per manufacturer’s protocol using luminometer (Promega Glomax 20/20) and data was analysed as stated in statistical analysis section. The lysates from the 3 triplicate wells were pulled together for western blot analysis.

**Figure 3:**
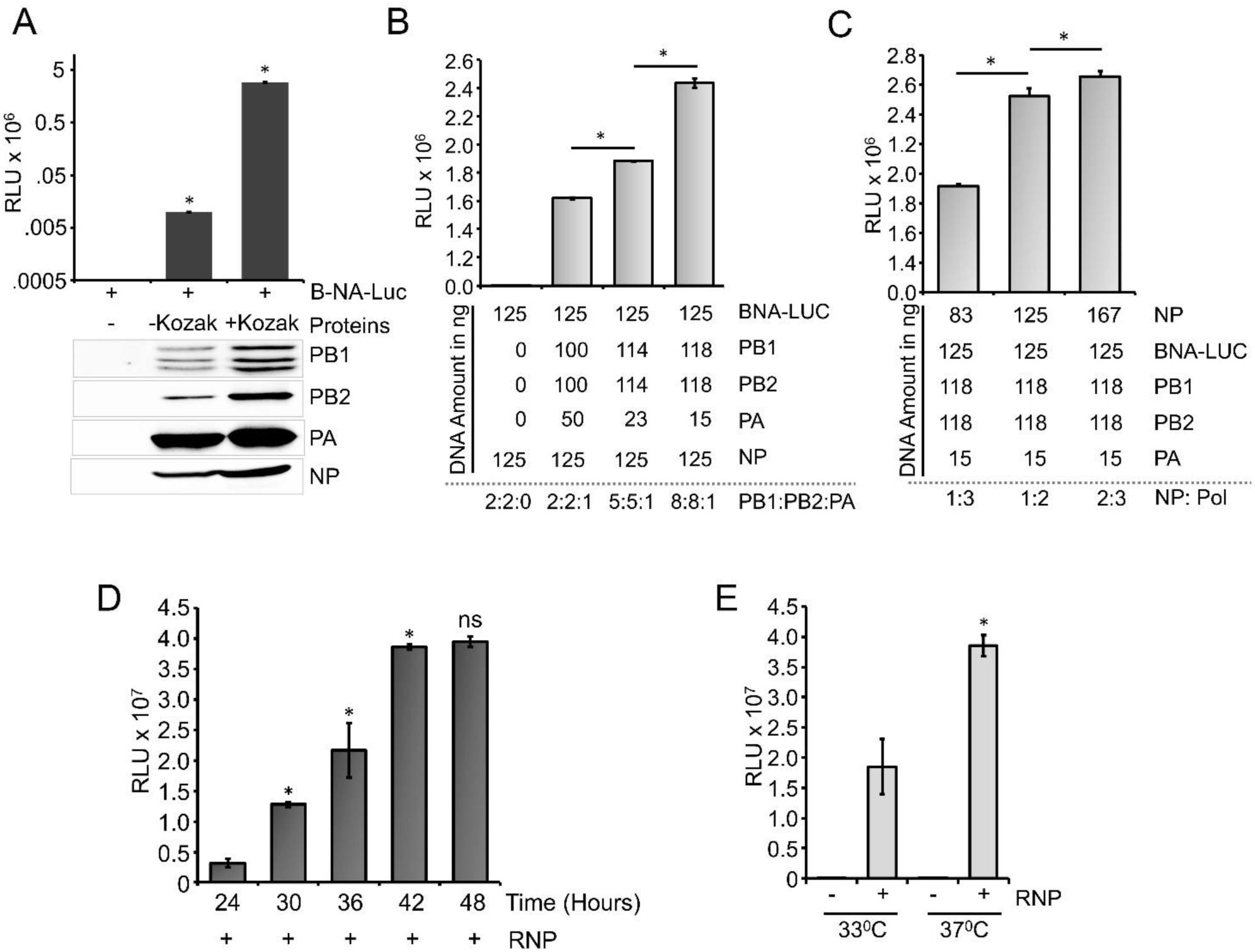
Optimization of the reporter system: (A) Effect of Kozak sequence on expression of viral polymerase proteins and influenza B RNP activity assay (B) Reporter RNP activity assay with different ratio of PA protein expression plasmid with respect to PB1 and PB2. (C) Reporter RNP activity assay with various amount of NP expression plasmid. (D) Optimization of time for reporter activity assay (E) Optimization of incubation temperature for influenza B RNP activity assay (n=3 ± standard deviation, *p<0.05 one-way ANOVA with post hoc Student’s t-test when compared to the preceding set, for Figure E, comparison was performed in between two RNP positive sets, ns = not significant).

**Figure 4:**
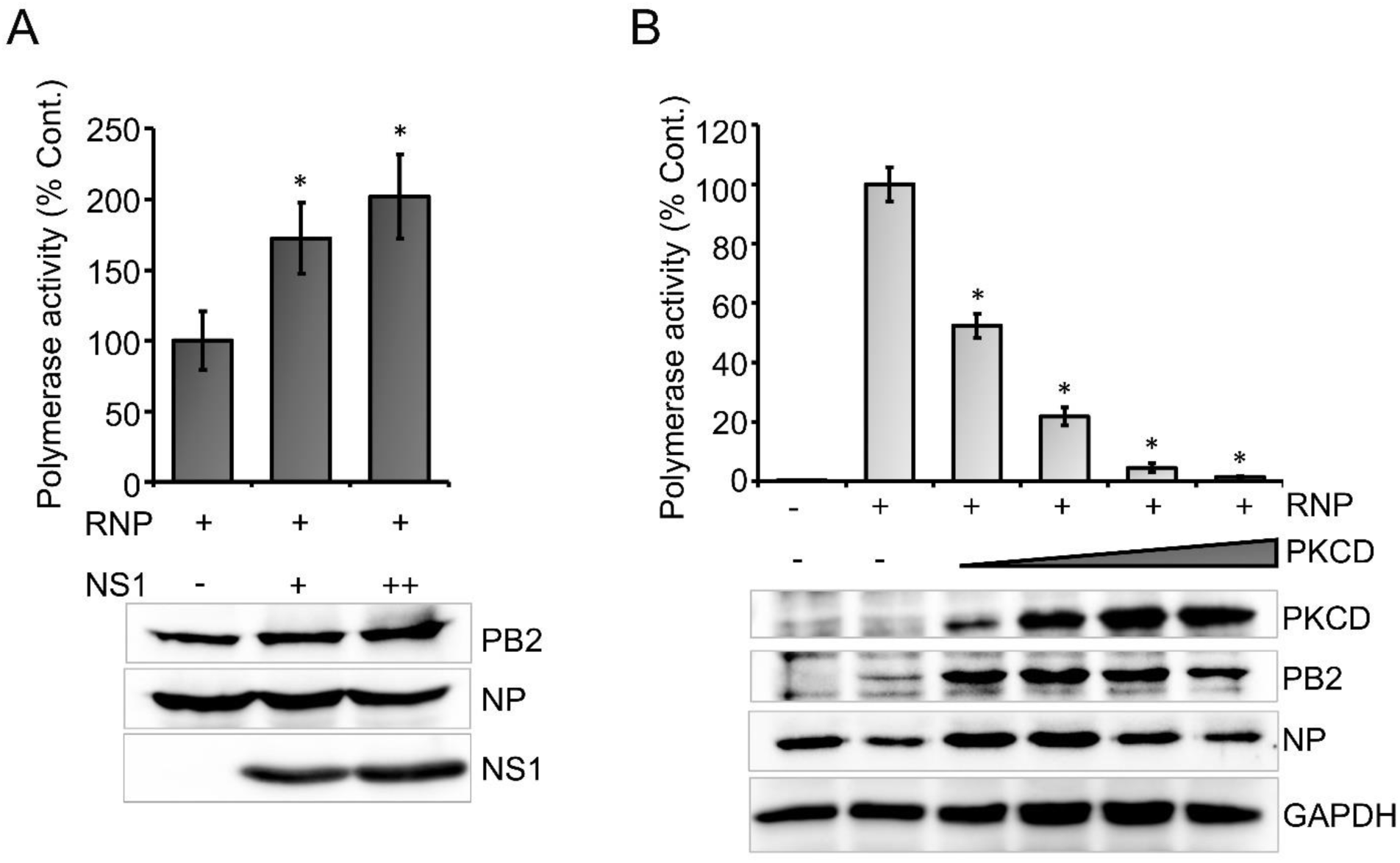
Effect of host & viral factors upon viral RNA synthesis: (A) Effect of an increasing amount of viral NS1 protein on Influenza B RNP activity assay (B) Effect of an increasing amount of constitutively active host protein kinase c delta (PKCD) protein on B RNP activity assay. (n=3± standard deviation. *p<0.05 one-way ANOVA with post hoc student’s t-test when compared to the preceding set.

**Figure 5:**
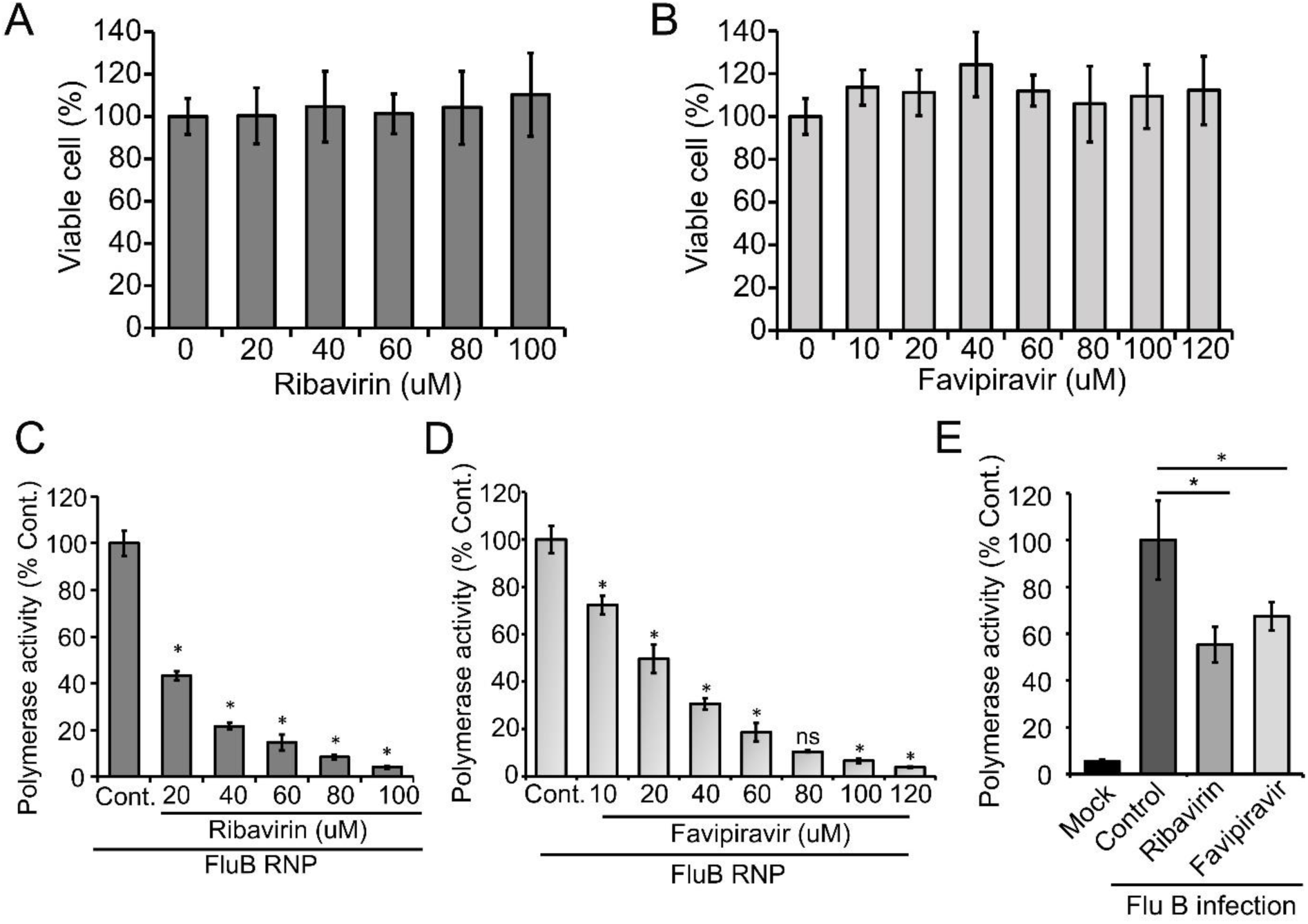
Effect of antiviral drugs upon viral RNA synthesis in infection-free and infection setting: (A, B) MTT assay to determine the cytotoxicity of Ribavirin and Favipiravir on HEK293T cells. (C, D & E) Effect of Ribavirin and Favipiravir on influenza B virus RNP activity. Viral polymerase proteins in HEK293T cells are expressed either by transient transfection (C, D) or by infecting the cells with Influenza B virus (E). (n=3 ± standard deviation, *p<0.05 one-way ANOVA with post hoc Student’s t-test when compared to the preceding set, for Figure E, comparison was performed with control set, ns = not significant).

### 2.10 Ribavirin and Favipiravir dose-response assays in HEK293T cells

Ribavirin (Sigma-Aldrich, Cat no. # R9644) was dissolved in water to prepare 80 mM stock, aliquoted and stored at – 20 ºC refrigerator. Favipiravir (MedChemExpress, Cat no. #HY-14768) was dissolved in DMSO (Sigma, Cat no. #D2650) to prepare a 200 mM stock, aliquoted and stored at – 80ºC until used. Working stocks for both the compounds were prepared in complete media. 0.2×10^6^ HEK293T cells were seeded in 24-well plates and post 24 hours were treated with specified concentrations of ribavirin or favipiravir for 2.5 hours at 37ºC with 5% CO2. Subsequently, cells were transfected with lipofectamine 3000 as per manufacturers protocol and incubated in fresh media containing specified concentrations of drugs were for 36 hours. Polymerase activity was then assayed as described above. The IC50 value was calculate by fitting the data to four parameter nonlinear equation.

### 2.11 3-(4,5-Dimethylthiazol-2-yl)-2,5-diphenyltetrazolium Bromide (MTT) Assay

3×10^4^ HEK293T cells were seeded in a 96-well plate. After 24 hours of seeding, the cells were treated with different concentrations of drugs in triplicates for 36 hours. Post-treatment, 100 μL of MTT reagent (5 mg/ml, SRL, Cat No. # 33611(2049101)) dissolved in phosphate-buffered saline (PBS) was added to the cells and incubated for 3 hours at 37ºC. Subsequently, the reagent was removed and the formazan crystals were dissolved by adding 100 μL of dimethyl sulfoxide (DMSO) to each well. The absorbance of the suspension was measured at 595 nm using an Epoch 2 microplate reader (BioTek Instruments). The percentages of drug treated metabolically active cells were compared with the percentage of control cells treated with vehicle control.

### 2.12 Transfection-Infection assay

0.1×10^6^ HEK293T cells were seeded in 48-well plates. After 20 hours of seeding, the cells were transfected with 0.25 ug of reporter plasmid using lipofectamine 3000 and 22 hours post-transfection, the cells were pre-treated with half-maximal inhibitory concentrations (IC50) of the drugs (Ribavirin:18.54 µM; Favipiravir:25.46 µM) for 2.5 hours at 37ºC with 5% CO2. Following 2.5 hours of treatment, the cells were infected with Influenza B virus at a MOI of 0.1 in presence of drugs and the polymerase activity was assayed at 16 hours of post-infection.

### 2.13 Statistical analysis

The arithmetic mean and standard deviation of the firefly luciferase signal were calculated from three biological replicates for each experiment. The mean intensity of luminescence (in arbitrary units) obtained from the luminometer was plotted in a bar diagram with standard deviations as error bars. For dual luciferase assay, the polymerase activity was reported as the ratio of firefly to renilla signal. In firefly luciferase assays involving the host factors, viral factors, antiviral molecules and dual luciferase assays; normalized mean and standard deviation were calculated against the control. The normalized mean was calculated by dividing the arithmetic mean of the experimental sets by the mean of the control set and converting them into a percentage value. Normalized standard deviation was calculated by normalizing the coefficient of variations against the control and augmenting it with the normal mean. A two-tailed Student’s T-test was performed for comparison of individual data sets. Intra assay variability was analysed by calculating the percentage coefficient of variation (%CV) of different sets with biological replicate. Student’s T-test was performed to compare the %CV of different assays. Inter-assay % CV was calculated from the mean %CV of three independent experiments.

## 3. Results

### 3.1 Generation of influenza B virus reporter plasmid for expression of viral reporter RNA in mammalian cells

In order to establish a reporter based RNP activity assay, a template RNA harboring the reporter gene flanked by the viral UTRs needs to be expressed under the control of RNA polymerase I promoter. This ensures that the reporter RNAs are devoid of any 5′- or 3′-terminal modifications, hence mimicking authentic viral genomic RNA. To achieve this, firstly we have constituted the “insert” harboring the firefly luciferase gene in reverse orientation flanked by viral 5′- and 3′-UTRs. Subsequently, this cassette was introduced into the pHH21 vector in between the RNA polymerase I promoter and terminator. The entire process of constituting authentic viral UTRs, assembling them with the reporter gene and introducing this cassette into the pHH21 vector utilized a single DNA polymerase enzyme without the need for any restriction enzyme or specialized kits (Figure 1). The sequences of 5′UTR, 3′UTR and primers corresponding to annealing regions are depicted in supplementary Figure S1.

Viral genomic RNA, purified from influenza B/Brisbane/60/2008 virus particles (as depicted in Figure 1A), was used as a source for the amplification of the 5′- and 3′-UTRs (103 nucleotides and 53 nucleotides respectively), using sequence specific primers with 18-24 nucleotide overhanging sequences corresponding to the vector and the reporter gene (Figure 1B, 2A). The resulting PCR products thus contain (i) viral 5′UTR flanked by the overlapping sequences with the Pol-I promoter and 3′-termini of the reporter gene of the total 144 base pairs in length and (ii) viral 3′UTR region flanked by the overlapping sequence with 5′-termini of reporter gene and Pol-I terminator of total 95 base pairs in length. These double-stranded PCR products were then used as primers to amplify the firefly luciferase gene from the pHH21-vNA-Luc plasmid, kindly provided by Prof. Andrew Mehle, University of Wisconsin Madison (Figure1C, 2B). The final PCR product, constitutes reporter gene flanked by viral 5′- and 3′-UTR regions along with partial sequences from the Pol-I promoter and terminator regions at the extreme 5′- and 3′-termini respectively. In order to synthesize the final reporter plasmid construct, named as pHH21-BNA-Luc, this cassette was inserted into the pHH21 vector (amplified in a separate PCR reaction; Figure 2C) using the Circular Polymerase Extension Cloning (CPEC), as originally described by Quan et al(32,34) (outlined in the Figure 1D). A vector to insert molar ratio of 1:3 generated maximum amount of assembled product (Figure 2D). Reaction product was transformed in chemically competent *E. coli* and the successful incorporation of the insert was confirmed by the colony PCR screening method. All the PCR amplifications were performed using a single Phusion High-Fidelity DNA polymerase as described in further detail in the methods section.

To reconstitute functional reporter RNPs inside the cells, reporter RNA template needs to be co-expressed with NP and the RdRp subunits, PB1, PB2 and PA (Figure 1E, F). The RdRp subunits were cloned into the pCDNA-3×-FLAG vector (generously provided by Dr. Andrew Mehle) under the control of CMV promoter with the help of EcoRI and NotI restriction enzymes, which results in incorporation of a tri-alanine linker in between the individual ORFs and the three FLAG epitopes joined in tandem (3×-FLAG). For expression of untagged proteins, ORFs with the stop codon were cloned using the same strategy. The NP gene was cloned into a modified pcDNA3 vector harboring V5 epitope tag (mentioned in the methods section) with the help of the KpnI and BamHI sites, with a glycine-glycine-serine-glycine linker in between the ORF and the V5 epitope tag.

### 3.2 Standardization of influenza B virus RNP activity assay

Influenza B virus reporter RNPs were reconstituted in HEK293T cells through transient transfection of the reporter plasmid (pHH21-BNA-Luc) either in the absence (negative control) or the presence of the plasmids expressing PB1, PB2, PA and NP proteins (Figure 3A). Cells were harvested at 24 hours of post-transfection and luciferase activity was measured to quantitate the Influenza B RNP activity. To our surprise, the positive control set showed only ∼10^4^ signal (luciferase light unit or RLU) which is only two log higher than the negative control set (∼10^2^ RLU), suggesting suboptimal activity of the reconstituted RNPs. This could be due to the poor expression levels of the RdRp subunits, PB1 and PB2, in comparison to the PA and NP proteins, as observed from the western blot analysis using specific antibodies, hence prohibiting the successful assembly of reporter RNPs inside the cells. We analysed the expression of individual polymerase proteins, PB2, PB1 and PA by transfecting them at increasing amounts into HEK293T cells and further western blot analysis as shown in supplementary Figure S2. It was observed that the expression of PA subunit is significantly higher than PB2 and PB1. To investigate this further, we examined the sequences of the constructs carefully and noticed that all of the protein expression plasmids lack the upstream Kozak sequence which may result in their suboptimal translation. The conserved Kozak sequence (GCCRCC**<u>ATG</u>**G) plays the critical role for recognition initiator ATG codon by ribosome to attain a high level of translation. Two specific positions, -3 and +4 from the adenine of initiator codon ATG are found (GCC**R**CC<u>ATG</u>**G**) to be critical for the optimal protein expression(35–38). The PA gene have a Guanosine at +4 position in its ORF but PB1 and PB2 do not have this Guanosine in their ORF. This makes the expression of PA gene better than PB2 and PB1. To address this, we performed site-directed mutagenesis to introduce partial Kozak sequences in each of these plasmids without any alteration in the ORF and repeated the polymerase activity assay with them. As evident from Figure 3A, the introduction of the Kozak sequence significantly boosted the expression of all of the RNP proteins which together resulted in reporter activity of 10^6^ RLU, four logs higher than the negative control set. Interestingly, the expression levels of the PA subunit still remained severalfold higher than the other two subunits of RdRp, PB1 and PB2 (Figure 3A). Precise abundance of the PB1, PB2 and PA subunits in equimolar amounts is a pre-requisite for the successful assembly of the heterotrimeric RdRp complex and hence, reconstitution of reporter RNPs to optimum levels. Therefore, we tried to optimize the amount of the plasmids to be transfected in order to have a comparable expression of the RdRp subunits. Reporter RNPs were reconstituted using different ratios of RdRp subunit plasmids, while keeping the amount of the reporter RNA and NP plasmids constant. As shown in Figure 3B, increasing the amount of PB1 and PB2 expressing plasmids compared to the PA led to a gradual increase in reporter activity and a ratio of 8:8:1 for PB1: PB2: PA resulted in comparable expression of all three polymerase subunits and maximum reporter activity. Protein expression level for all polymerase subunits have been shown by western blot analysis in supplementary Figure S3. Subsequently, keeping the ratio of the polymerase subunit plasmids constant, we increased the amount of the NP expressing plasmid, which resulted in increase in the reporter activity, hence stretching the sensitivity of this reporter assay to the maximum level (Figure 3C). Individual protein expression levels are shown in supplementary Figure S4 as western blot analysis. The NP to polymerase proportion up to 1:2 results in increase in polymerase activity. Further increase in the amount of NP does not result in substantial increase in polymerase activity. Hence for our further experiments, we have used this ratio of RNP reconstituting plasmids. Once we optimize the amounts of various plasmids reconstituting reporter RNPs, we have performed a time kinetics experiment in order to assess the optimum time required to obtain a signal of 10^6^ RLU or more (Figure 3D).

A time-dependent increase in the reporter activity was observed which reaches a plateau at 42 h post transfection. Additionally, influenza B polymerase activity was assessed at different temperatures (33 ºC and 37 ºC) by reconstituting the polymerase through transient transfection at 37 ºC for 12 hours followed by an additional incubation of 30 hours at respective temperatures (Figure 3E). As observed at 37 ºC, reporter activity was almost two fold higher than the activity at 33 ºC, a data corroborated perfectly with the previous results obtained by Santos et al(39). Together, we present a fast sensitive and high throughput reporter assay for monitoring influenza B virus RNA synthesis in an infection free setting.

All of the assays were performed in 24-, 48- and 96-well plates, in triplicates for each of the biological sets (data presented in this manuscript is from 24 well plates), hence confirming that this assay system is high-throughput compatible. Additionally, five log difference between the signal and background readouts provides a wide dynamic range for this reporter-based assay system. There is no significant difference in the intra-assay variability among individual experiments as shown in supplementary Figure S5. The inter-assay coefficient of variation remains within 10% (supplementary Figure S5). Together, we have been able to establish a fast, reliable and high-throughput compatible assay system for monitoring influenza B virus RNP/ polymerase activity, which is suitable for assessing the effect of various viral or cellular factors in modulating RNP activity and hence viral RNA synthesis.

### 3.3 The influenza B RNP activity assay is suitable for evaluating the efficacy of viral or host factors in regulating viral RNA synthesis

To this end, we set out to evaluate the efficacy of the newly developed polymerase activity assay in identifying novel viral and host factors that may regulate viral RNA synthesis. Influenza virus Nonstructural Protein 1 (NS1) is a multifunctional protein participating mainly in the suppression of antiviral defense mechanisms exerted by a wide variety of host factors (40,41). Additionally, influenza A virus NS1 protein has been shown to boost viral RNA synthesis(42–44), possibly through interfering with antiviral activity of DDX21 and RAP55 (45,46). While the immune suppression activity of influenza B NS1 was well studied(41,47,48), little is known about the role of NS1 in regulating influenza B virus RNA synthesis. Hence, we evaluated the ability of influenza B virus NS1 protein to promote viral RNA synthesis with the newly developed reporter RNP activity assay. Influenza B virus NS1 ORF was cloned into the pCDNA-3×-FLAG vector that resulted in the expression of the C-terminally FLAG tagged NS1 protein. Influenza B reporter RNPs were reconstituted in HEK293T cells either in the absence or presence of increasing amounts of NS1 protein and reporter activity was monitored to assess the extent of viral RNA synthesis. Increasing amount of NS1 resulted in 1.5 to 2 folds increase in reporter activity (Figure 4A) establishing it as a positive regulator of viral RNA synthesis. Furthermore, reconstituting reporter RNPs in the presence of NS1 presents an assay system that closely resembles RNA synthesis, which occurs during the course of infection.

Subsequently, we tested the ability of a host factor to regulate influenza B virus RNA synthesis using our reporter RNP activity assay. Host Protein Kinase C, specifically the delta isoform, has been shown to positively influence influenza A virus RNA synthesis by regulating the phosphorylation and subsequent assembly of viral nucleoproteins into RNPs. Interestingly, the constitutively active catalytic domain of PKC delta (PKCD), when overexpressed, negatively regulates influenza A virus RNA synthesis (49). To determine the role of PKCD in regulating influenza B virus RNA synthesis we employed the newly developed reporter RNP activity assay. As evident from Figure 4B, increasing amounts of PKCD resulted in a gradual decrease in RNP activity and hence viral RNA synthesis without any severe impact upon the translation of viral proteins. These data not only substantiate the role of PKCD in regulating influenza B virus RNP activity, but also validates the efficacy of our assay system in studying the effect of pro- or antiviral factors regulating viral RNA synthesis.

### 3.4 Reporter based RNP activity assay as high throughput screening platform for antiviral drugs

Finally, we intend to establish the suitability of the RNP activity assay as a high throughput screening platform for antiviral drugs that can inhibit viral RNA synthesis and hence virus replication. Ribavirin and Favipiravir are nucleoside (purine) analogues, which inhibit the replication of a wide variety of RNA viruses by acting as an alternative substrate for viral RNA polymerase(50,51). Additionally, Ribavirin also inhibits inosine monophosphate dehydrogenase thereby depleting the GTP and creating an imbalance in the nucleotide pool inside the cell(52). Both Ribavirin and Favipiravir has been approved as chemoprophylaxis as well as therapy against influenza A and B viruses(53–55). Hence, we used these two drugs as positive controls to test the efficacy of our assay system for antiviral screening. MTT assay was performed in HEK293T cells (Figure 5A, B), where neither of the drugs show any cytotoxicity upto investigated concentrations. HEK293T cells were pretreated with different concentrations of the drugs followed by forward transfection to reconstitute the influenza B reporter RNPs and subsequent incubation with the drugs for 36 hours. Reporter activities were measured and expressed as relative percentages with respect to the vehicle control. Data presented in Figure 5 C, D, shows a dose-dependent decrease in the reporter activity and hence viral RNA synthesis with increasing amounts of the drugs with IC50 values of 18.54 µM and 25.46 µM for Ribavirin and Favipiravir respectively.

To further extend the scope of the assay system, we sought to check if this system is capable of assessing the effect of host factors or antivirals upon the overall progress of infection. For this purpose, we transfected HEK293T cells with the reporter construct and subsequently infected them with influenza B virus at 20 hours post transfection. It is expected that in infected cells reporter RNA template will get transcribed with the help of RdRp and NP proteins expressed from viral genomic RNA segments. As evident from Figure 5E, infected cells supported successful generation of reporter RNPs and hence showed high reporter activity, while the uninfected cells showed no such effect. Interestingly, when parallel set of cells were treated with Ribavirin and Favipiravir prior to infection with Influenza B virus, significant reduction in reporter activity were observed, hence suggesting an overall reduction in viral gene expression and hence virus replication in presence of the drugs. Together our data reconfirms the activity of the two well established antiviral drugs against influenza B virus RNA synthesis machinery and also establishes the newly developed reporter-based influenza B virus RNP activity assay as a high throughput screening platform for antivirals specifically inhibiting viral RNA synthesis.

## 4 Discussion

Luciferase based reporter assay systems remain a key tool for analyzing gene expression in a wide variety of organisms; viruses are not exception. While, for positive sense RNA viruses, introduction of the single sub-genomic reporter RNA template in cells is sufficient for expression of reporter genes; but for negative sense RNA viruses, RNP associated viral proteins needs to be synthesized along with the reporter RNA in order to reconstitute complete RNPs, which then leads to the expression of reporter enzyme as a proxy of viral genes(20,56,57). This is why, successful reconstruction of reporter viral RNPs require extensive cloning of multiple RNA and protein expressing constructs, standardization of their expression in right stoichiometric ratios and optimization of other crucial parameters like time, temperatures etc. Although, several groups have reported reporter assay systems for monitoring influenza A and B virus RNA synthesis, non-availability of detailed methodical description makes the process of establishing the assay system non-trivial(13,20,25–27). In this work, we have established a firefly luciferase-based influenza B virus RNP activity assay and presented the detailed methodology of the entire procedure which could be easily followed for the development of such viral and non-viral reporter assay systems.

We have introduced a unique cloning strategy for the construction of the influenza B virus reporter RNA construct that is devoid of restriction enzymes or any other specialized enzymes. This cloning strategy utilizes a single DNA polymerase, which is widely used for regular molecular biology work and hence easily available. Using this polymerase, two consecutive PCR amplification reactions led to the generation of the reporter RNA cassette encompassing the reporter ORF flanked by viral 5′- and 3′-UTR regions which were then inserted into the vector using CPEC cloning method. While the vector and the insert used for CPEC, are also compatible for Gibson assembly based cloning method, we intentionally avoided use of any specialized enzymes to make the overall procedure simple and user-friendly that could be adapted for cloning of any other reporter RNA constructs. In addition to reporter RNA construct, we also cloned ORFs corresponding to viral PB1, PB2, PA and NP proteins and optimized their expression to reconstitute reporter RNPs at maximum levels. The robustness of this assay system was substantiated by testing the efficacy of antiviral drugs, Ribavirin and Favipiravir, to inhibit influenza B virus RNA synthesis either in the context of reconstituted RNPs (through transfection) or during the course of infection. The fact that the reporter RNA template can be preferentially recognized by viral NP and RdRp subunits to reconstitute reporter RNPs during the course of infection, confirms that the reporter RNA mimics viral genomic RNA segments and hence validates its suitability to be used for the study of viral RNA synthesis and effect of various viral and host factors upon the same. In fact, using the newly developed reporter RNP system, we for the first time showed that viral NS1 protein can boost influenza B virus RNA synthesis and constitutively active form of host PKCD can downregulate the same. While effect of NS1 and PKCD proteins has been previously characterized in case of influenza A virus(42–44,49), our results substantiates that these proteins participate similarly regulate influenza B virus RNA synthesis as well.

Altogether, we present a comprehensive roadmap for development, characterization and validation of a reporter-based Influenza B virus polymerase/ RNP activity assay and made it generic enough to be followed by others who intend to develop similar assay systems for influenza and other negative sense RNA viruses. We also made all the resources publicly available (upon request) to enrich the armoury for combating influenza B viruses and hope that it will be widely utilized to identify new therapeutic strategies against this deadly human pathogen.

## Supporting information

https://www.frontiersin.org/articles/10.3389/fmicb.2022.868367/full#supplementary-material

## 6 Author Contributions

N.K designed and performed the experiment, standardized methodology, analysed the data, wrote the original draft, reviewed and edited the manuscript. S.B. - designed and performed experiments, analyzed the data, reviewed and edited the manuscript. A.M conceptualized the work, designed and supervised the project, arranged for funds, wrote the original draft, reviewed and edited the manuscript. All authors have read and approved this final manuscript.

## 8 Acknowledgments

We sincerely acknowledge Prof. Andrew Mehle (University of Wisconsin Madison) for proving us various valuable plasmids. We acknowledge Indian Institute of Technology Kharagpur for providing us infrastructural support. A.M. acknowledges DBT, Ramalingaswami re-entry fellowship (BT/RLF/Re-entry/02/2015), DST-SERB, Early Career Research Award (ECR/2017/001896) and MHRD, “Scheme for Transformational and Advanced Research in Science” {STARS/APR2019/BS/369/FS (Project ID: 369)} for financial support. N.K. (File number 09/081(1316)/2017-EMR-I) and S.B. (File number 09/081(1301)/2017-EMR-I) acknowledge the Council of Scientific and Industrial Research for their fellowship.

## 1. Data Availability Statement

All data generated or analysed during this study are included in this published article.

