## Supplementary material for "A Comprehensive Roadmap Towards the Generation of an Influenza B Reporter Assay Using a Single DNA Polymerase-Based Cloning of the Reporter RNA Construct": https://www.frontiersin.org/articles/10.3389/fmicb.2022.868367/full#supplementary-material

This section contains supplementary figures S1-S5.

### Supplementary Figures

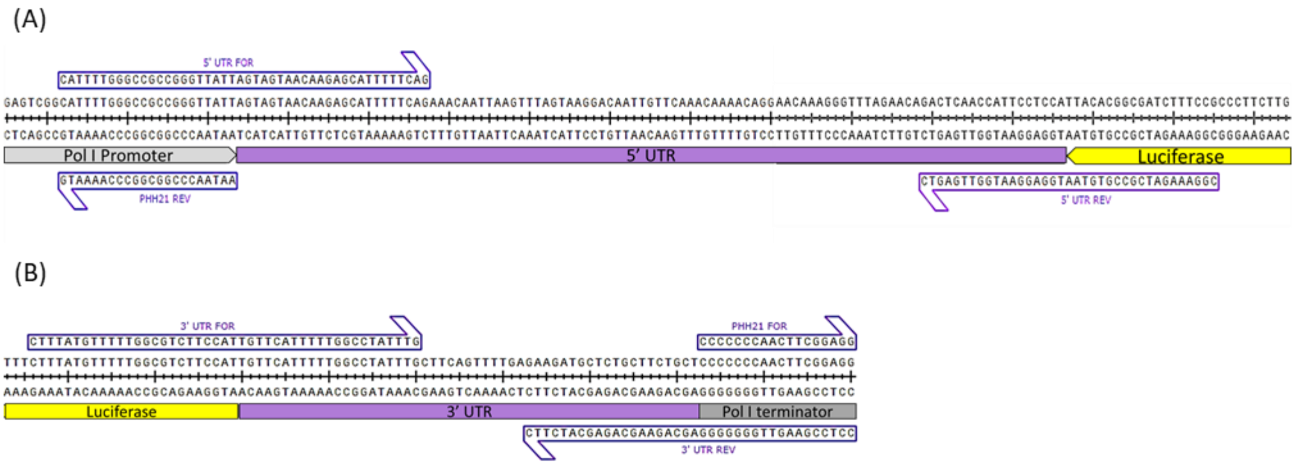


**Figure S1**: Schematic diagram showing alignment of primers with the 5’**(A)** and 3’**(B)** UTRs of influenza B virus segment 6 with overhangs aligning with luciferase ORF (in antisense orientation) and Human RNA polymerase-I promoter and terminator sequences.

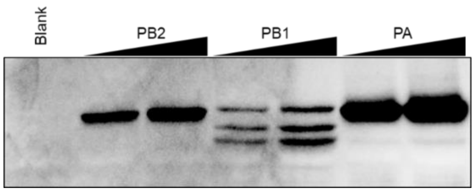


**Figure S2**: **Individual Expression of polymerase subunits in HEK293T cells**. Equally increasing amounts (250 ng and 500 ng) of mammalian expression plasmids were transfected in $2.5\times10^5$ HEK293T cells. 24 hours post transfection, cells were lysed and western blot was performed.

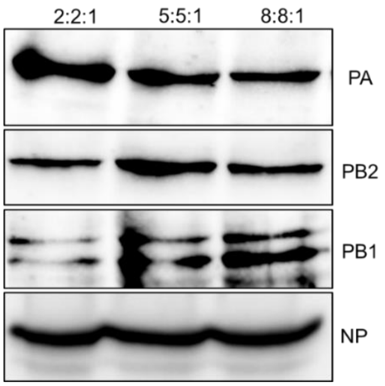


**Figure S3**: **Optimization of the polymerase subunit expression for efficient reporter activity.** Reporter RNP activity assay with different ratio of PA protein expression plasmid with respect to PB1 and PB2 was performed into the HEK293T cells (corresponding to Figure 3B). The cells lysates were subjected to western blot analysis. The ratio of PB2:PB1:PA respectively are mentioned above the lanes.

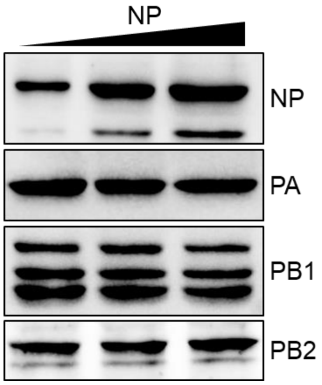


**Figure S4: Optimization of the nucleoprotein (NP) expression for efficient reporter activity.** Reporter RNP activity assay with increasing amount of NP expression plasmid was performed into the HEK293T cells (corresponding to Figure 3C). Western blots showing the expression of different polymerase sub-units and NP.

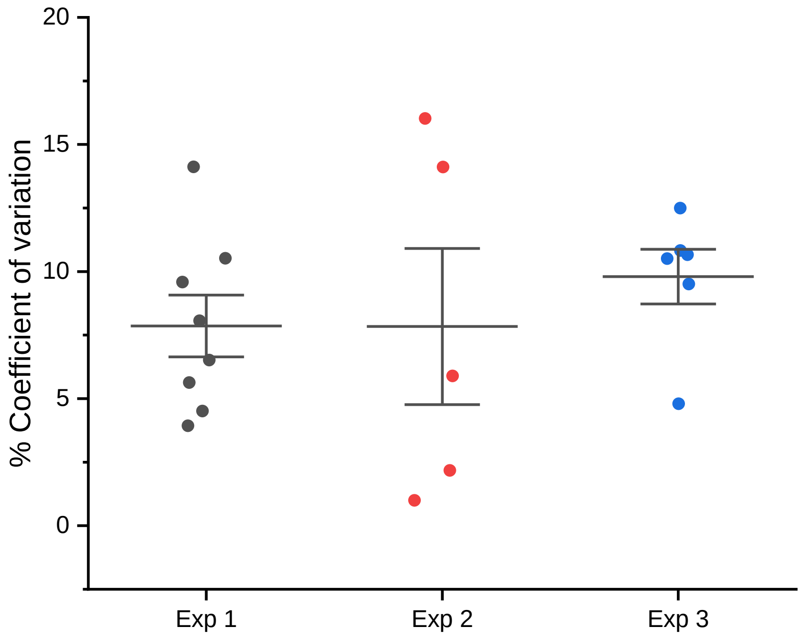


**Figure S5**: **Percent coefficient of variation (% CV) calculated from individual experiments to investigate the intra-assay and inter-assay variability**. Each dot represents %CV from three replicates for each set in one experiment. Middle bar represents the mean % CV ± standard error of one experiment. %CV of each experiments were compared with each other by student's t-test. No statistically significant difference was found among the experiments.
